# Accounting for heterogeneity due to environmental sources in meta-analysis of genome-wide association studies

**DOI:** 10.1101/2024.05.17.594687

**Authors:** Siru Wang, Oyesola O. Ojewunmi, Abram Kamiza, Michele Ramsay, Andrew P Morris, Tinashe Chikowore, Segun Fatumo, Jennifer L Asimit

## Abstract

Meta-analysis of genome-wide association studies (GWAS) across diverse populations offers power gains to identify loci associated with complex traits and diseases. Often heterogeneity in effect sizes across populations will be correlated with genetic ancestry and environmental exposures (e.g. lifestyle factors). We present an environment-adjusted meta-regression model (env-MR-MEGA) to detect genetic associations by adjusting for and quantifying environmental and ancestral heterogeneity between populations. In simulations, env-MR-MEGA had similar or greater association power than MR-MEGA, with notable gains when the environmental factor had a greater correlation with the trait than ancestry. In our analysis of low-density lipoprotein cholesterol in ∼19,000 individuals across twelve sex-stratified GWAS from Africa, adjusting for sex, BMI, and urban status, we identified additional heterogeneity beyond ancestral effects for nine variants. Env-MR-MEGA provides an approach to account for environmental effects using summary-level data, making it a useful tool for meta-analyses without the need to share individual-level data.

## Introduction

Genome-wide association studies (GWAS) have identified thousands of loci contributing genetic effects to a wide range of complex traits and diseases^1–4^. Although GWAS have successfully identified hundreds of genetic variants associated with complex diseases, those genetic variants explain a small proportion of heritability ^5,6^; many variants with smaller effect sizes would explain the remaining contribution. To identify these novel variants with modest effects, meta-analysis is the common approach to collect the large sample sizes needed across independent GWAS, without the need to exchange individual-level genotype and phenotype data.

When conducting a meta-analysis, it is expected that multiple GWAS are collected from highly similar studies to minimise heterogeneity in allelic effect of the same genetic variants across these GWAS^7,8^. Under this assumption, some fixed-effects models were developed^9–12^. However, even when these collected studies are derived from the same genetic ancestry, heterogeneity in the effect sizes across these GWAS would inevitably occur due to genetic ancestry, as well as environmental exposures that may interact with genetic variants; the fixed-effect analysis cannot accommodate this heterogeneity issue^13^. Some causes of allelic heterogeneity include variations in patterns of linkage disequilibrium (LD) between cohorts and varied environmental exposures across cohorts that could interact with the genetic contribution^14–18^.

Another approach for addressing the heterogeneous effect sizes is to meta-analyse with a random-effects model^19–21^. These traditional random-effects models estimate the effect sizes and their standard errors by assuming the effect sizes from the studies follow the same distribution^22^. It has been demonstrated that traditional random-effects models have lower power to detect associations than fixed-effects models at variants with varying effect sizes across cohorts ^23–25^. However, an alternative random-effects model ^25^ has been shown to have power gains over fixed-effects models by assuming there is no heterogeneity under the null hypothesis. Nevertheless, these approaches do not allow for the expectation that the extent of heterogeneity between a pair of GWAS is likely to be correlated with the genetic distance between them. In other words, we would expect structure for heterogeneity, whereby GWAS from the same ancestry group will be more homogenous than those from different ancestry groups.

To account for the heterogeneity structure between ancestry groups, MANTRA utilised a Bayesian framework to cluster different cohorts by a prior model of genetic similarity, which is assessed by mean pairwise genome-wide allele frequency differences^26^. Compared to the fixed-effects and random-effects meta-analysis, MANTRA has shown significant improvement in the power to detect associations when the similarity in allelic effects between populations is well captured by their relatedness^27^. However, MANTRA implements a Metropolis-Hastings MCMC algorithm ^28,29^, which would lead to a heavy computational burden for the meta-analysis of many GWAS. To address this computational challenge, MR-MEGA^30^ was developed, which in addition to detecting genetic associations, quantifies the heterogeneity in allelic effects that is due to genetic ancestry. MR-MEGA models allelic effects, weighted by the corresponding standard errors, as a function of axes of genetic variation derived from mean pairwise genome-wide allele frequency differences between GWAS. Compared to fixed and random-effects meta-analysis, MR-MEGA has increased power to detect association when there is heterogeneity in allelic effects between populations due to genetic ancestry^30^.

Until now, the MR-MEGA approach only accounts for the heterogeneity in allelic effects that is correlated with genetic ancestry. Increasing evidence shows that different environmental exposures between GWAS may also contribute to varying allelic effects across populations. For example, sex can be treated as a simple “environmental” risk factor, incorporating physiological and behavioural differences between males and females at different stages that may lead to differences in allelic effects between sexes ^31^. A series of examples of sex-differentiated effects were confirmed through human GWAS ^32–34^, demonstrating that sex can interact with causal variants, which causes the heterogeneity in allelic effects between males and females.

Considering the impact of environmental exposures that differ across GWAS, we have developed a novel environment-adjusted meta-regression that accounts for environmental exposures specific to each cohort, in addition to ancestry. Building on the MR-MEGA meta-regression framework, we construct the environment-adjusted meta-regression model, env-MR-MEGA, by adding study-level environmental covariates. This allows us to identify genetic variants that are associated with the disease or trait while adjusting for differing environmental exposures between cohorts.

In our extensive simulation study, we assessed the power to detect association and heterogeneity of allelic effects due to ancestry and/or environment using env-MR-MEGA. In our application to twelve sex-stratified cohorts of ∼19,000 individuals from Africa (West, East and South Africa), env-MR-MEGA identified nine variants for LDL cholesterol that have heterogeneity beyond that explained by ancestral effects.

## Results

Taking account of different environmental exposures between cohorts, we developed an environment-adjusted meta-regression model of GWAS in which the MR-MEGA meta-regression model framework ^30^ is adapted by adding study-level environmental covariates (Methods). This adjustment accounts for not only the heterogeneity in allelic effects that are correlated with genetic ancestry but also the heterogeneity in varied environmental exposures between GWAS. Based on the environment-adjusted meta-regression model, env-MR-MEGA, we test for genetic associations and the heterogeneity in allelic effects due to genetic ancestry and environmental effects or individually from each of genetic ancestry and environment.

In a series of conducted simulations, we obtained allele frequencies from eight African populations: Durban Diabetes Study (DDS) and Durban Case Control Study (DCC)^35^, which are two Zulu cohorts, the Uganda Genome Resource (UGD)^36^ and Esan (ESN), Gambian (GWD), Luhya (LWK), Mende (MSL), and Yoruba (YRI) from Phase 3 of 1000 Genomes^37^. Based on the variants with minor allele frequencies (MAF) >5% in all eight populations, we derived two axes of genetic variation, which suggest that the populations separate into four clusters: (i) ESN and YRI; (ii) GWD and MSL; (iii) UGD and LWK; (iv) DCC and DDS (**Supplementary Material FigS1**). These four clusters are consistent with geographical regions: (i) West-central Africa; (ii) West Africa; (iii) East Africa; (iv) South Africa. Then we incorporated the four clusters and introduced six heterogeneity scenarios for the presence of a genetic association: ancestrally homogeneous, East Africa, West Africa, West-central Africa, South Africa and non-ancestral Africa (**Supplementary Material Table S1**). For example, under the East Africa setting, populations from East Africa (UGD and LWK) are associated with a particular genetic variant, and all other populations are null at that variant.

Subsequently, a range of sex-stratified models of association across these heterogeneity scenarios were explored in simulations of 16 sex-stratified GWAS. Consequently, we assess the performance of env-MR-MEGA compared to MR-MEGA in a series of simulations of sex-stratified cohorts with varied sample sizes (equal medium-sized vs. unequal large-sized) and varied environmental exposure proportions across cohorts. In the application of env-MR-MEGA, we apply env-MR-MEGA to LDL-cholesterol in 12 sex-stratified cohorts from Africa, with environmental covariates sex, BMI, and urban status. We evaluate and assess evidence of heterogeneity due to these environmental sources and from ancestry.

### Adjusting for a stratified environment improves power in detecting genetic associations and allelic heterogeneity

Treating sex as an environmental covariate, based on 16 sex-stratified cohorts, we examined a model of female-specific association with the trait for the six heterogeneity scenarios (**Supplementary Material**, **Table S1**). As a baseline comparison, we consider equal-sized samples of size 1000, as well as varying sample sizes of 3000∼4000 in each cohort.

Across the six heterogeneity scenarios, type I errors for detecting associations are well-calibrated at 0.05 (**Supplementary Material**, **Table S4**). Among the six heterogeneity scenarios, under a model of female-specific effects, env-MR-MEGA gained the greatest power to detect association over MR-MEGA regardless of sample size **(Fig 1**). Between different sample size settings in each cohort, smaller sample sizes in female/male cohorts (1000) have substantially lower power than the larger mixed sample sizes (≥3000 in each cohort), as expected. For scenarios in which heterogeneity in female allelic effects was correlated with ancestry (East Africa, West-central Africa, West Africa and South Africa), no matter whether the sample size in each cohort is equal or not, env-MR-MEGA provided the greatest power to detect association. For the scenarios in which heterogeneity in female allelic effects across populations was homogeneous and random (non-ancestral Africa), env-MR-MEGA still attained the greatest power, as expected. In addition, compared with the above-mentioned four scenarios in which female allelic effects varied across African regions (East Africa, West-central Africa, West Africa and South Africa), the gains in power over MR-MEGA were higher.

**Figure 1.**
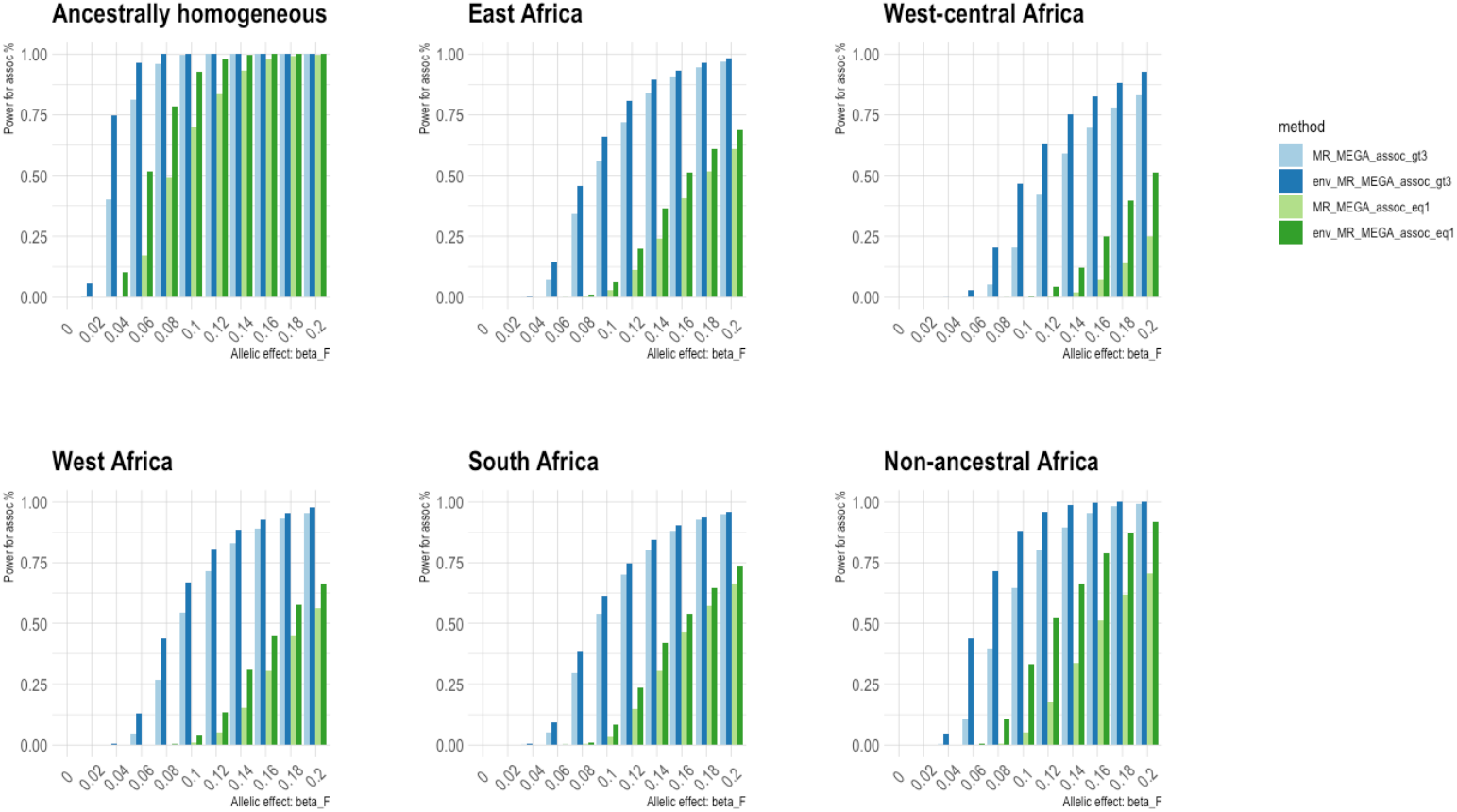
Power to detect association by env-MR-MEGA is higher than MR-MEGA in various heterogeneity scenarios involving 16 sex-stratified cohorts from east, west, west-central and south African regions. “MR_MEGA_assoc_gt3” and “env_MR_MEGA_assoc_gt3” correspond to the setting of unequal sample sizes (≥3000 in each cohort); **“**MR_MEGA_assoc_eq1” and “env_MR_MEGA_assoc_eq1” correspond to the setting of equal sample sizes (1000 in each cohort). Power was assessed at *P* < 5 × 10^−8^ and based on 1000 simulations.

For a more comprehensive exploration of the ancestrally homogeneous scenario, we extended the homogeneity in ancestry alone to homogeneity in ancestry and sex, where both female and male cohorts in each African population share the same allelic effects instead of female cohorts only. Tests for heterogeneity due to ancestry and environment by env-MR-MEGA are well-calibrated (**Supplementary Material**, **Table S5**). In the model of homogeneity of allelic effects in ancestry and sex between males and females, the type I errors for detecting (i) heterogeneity due to ancestry and environment was 0.048 (0.049 under the equal-sized setting); (ii) heterogeneity due to ancestry was 0.054 (0.040 under the equal-sized setting); (iii) heterogeneity due to environment was 0.045 (0.053 under the equal-sized setting).

Among the heterogeneity scenarios of East Africa, west-central Africa, west Africa and South Africa, the power for heterogeneity in allelic effects due to the ancestry and environment obtained from env-MR-MEGA was greatest (**Supplementary Material**, **Fig S2**). As expected, the power to detect allelic heterogeneity due to ancestry was the same for env-MR-MEGA and MR-MEGA. The two settings of unequal sample size and equal sample size gave similar results for the heterogeneity tests, with the expected lower power among the smaller sample tests (**Supplementary Material**, **Fig S3**). Furthermore, the power for heterogeneity due to environment was greater than that due to ancestry in the west-central Africa scenario while the power for heterogeneity due to environment was lower than that due to ancestry in the other ancestry-specific heterogeneity scenarios (east Africa, west Africa and south Africa). This suggests that the allelic heterogeneity in the west-central Africa scenario shows lower correlations with ancestry compared to sex effect, unlike in other ancestry-specific scenarios, East Africa, West Africa and South Africa, where allelic heterogeneity shows a stronger correlation with ancestry. Possibly this is because the axes of genetic variation for west-central Africa are between those for West Africa and East Africa (**Supplementary Material**, **Figure S1**).

For the ancestrally homogeneous scenario, as expected, the power to detect allelic heterogeneity due to ancestry from both env-MR-MEGA and MR-MEGA attained the nominal significance threshold, α = 0.05 (**Supplementary Material**, **FigS2 and FigS3**). Specifically, the power for allelic heterogeneity due to environment was slightly higher than the power for allelic heterogeneity due to ancestry and environmental effects. In summary, the power for allelic heterogeneity involving environmental effects was notably greater than the power for heterogeneity due to ancestry alone. This is because the heterogeneity in allelic effects across sex-stratified cohorts was correlated with sex alone. For the non-ancestral Africa scenario, the power to detect heterogeneity was noticeably lower for ancestry effects only compared to environmental effects because of the randomness in assigned effects among population groups.

### Adjusting for environment proportions improves power in detecting genetic associations and allelic heterogeneity

To investigate the impact of the environmental effect, we considered a more complicated setting in which the heterogeneity in allelic effect is correlated with the environmental factor instead of sex alone. Here, smoking status was treated as the environmental covariate, and the proportion of smokers varied across the eight populations, including the sex-stratified 16 male/female cohorts. The smoking proportions can vary not only across male and female cohorts but also across different ancestries. To explore a broader range of scenarios, we considered two smoking patterns: (i) small differences in smoking proportions between males and females but large differences in smoking proportions between populations in the same African region (**Supplementary Material Table S2**); (ii) large differences in smoking proportions between male and females but minor differences in smoking proportions between populations in the same African region (**Supplementary Material Table S3**). Additionally, for each smoking pattern, we considered three settings: (a) no difference: female and male cohorts in the same population shared the same smoking proportions; (b) same direction: smoking proportions in female cohorts were always lower than that in male cohorts; (c) mixed direction: among the eight populations, not all female smoking proportions were lower than that in male cohorts. Consequently, through a range of conducted simulations under these settings, it is demonstrated that the env-MR-MEGA had the highest gains in power over MR-MEGA when there was higher variability in smoking proportions between the cohorts.

In the first smoking pattern, where male and female cohorts from the same population share identical smoking proportions or varied smoking proportions with small differences (**Supplementary Material Table S2**), env-MR-MEGA gained the greatest power to detect association over MR-MEGA, particularly in ancestrally homogeneous, west-central Africa and non-ancestral Africa scenarios (**Fig 2, Supplementary Material FigS4**, **FigS5**). For East Africa and West Africa scenarios, env-MR-MEGA gained slightly higher power than MR-MEGA, whereas, in the South Africa scenario, the power to detect association was nearly equivalent for both env-MR-MEGA and MR-MEGA. It can be explained that when heterogeneity in allelic effects is more strongly correlated with the ancestry compared to smoking exposures, especially in the South Africa scenario where the two Zulu cohorts, DCC and DDS, are highly similar to each other (**Supplementary Material**, **FigS1**), the power for association would be more significantly influenced by ancestry compared to smoking exposures. Consequently, this results in a similarity in power for the association between env-MR-MEGA and MR-MEGA in the South Africa scenario. Additionally, under this smoking pattern, there was no notable difference in power of association in each heterogeneity scenario between these three settings for the varied directions of difference in smoking proportions between males and females, which include no difference, mixed direction and same-direction (**Supplementary Material**, **Fig S4**, **Fig S5**).

**Figure 2.**
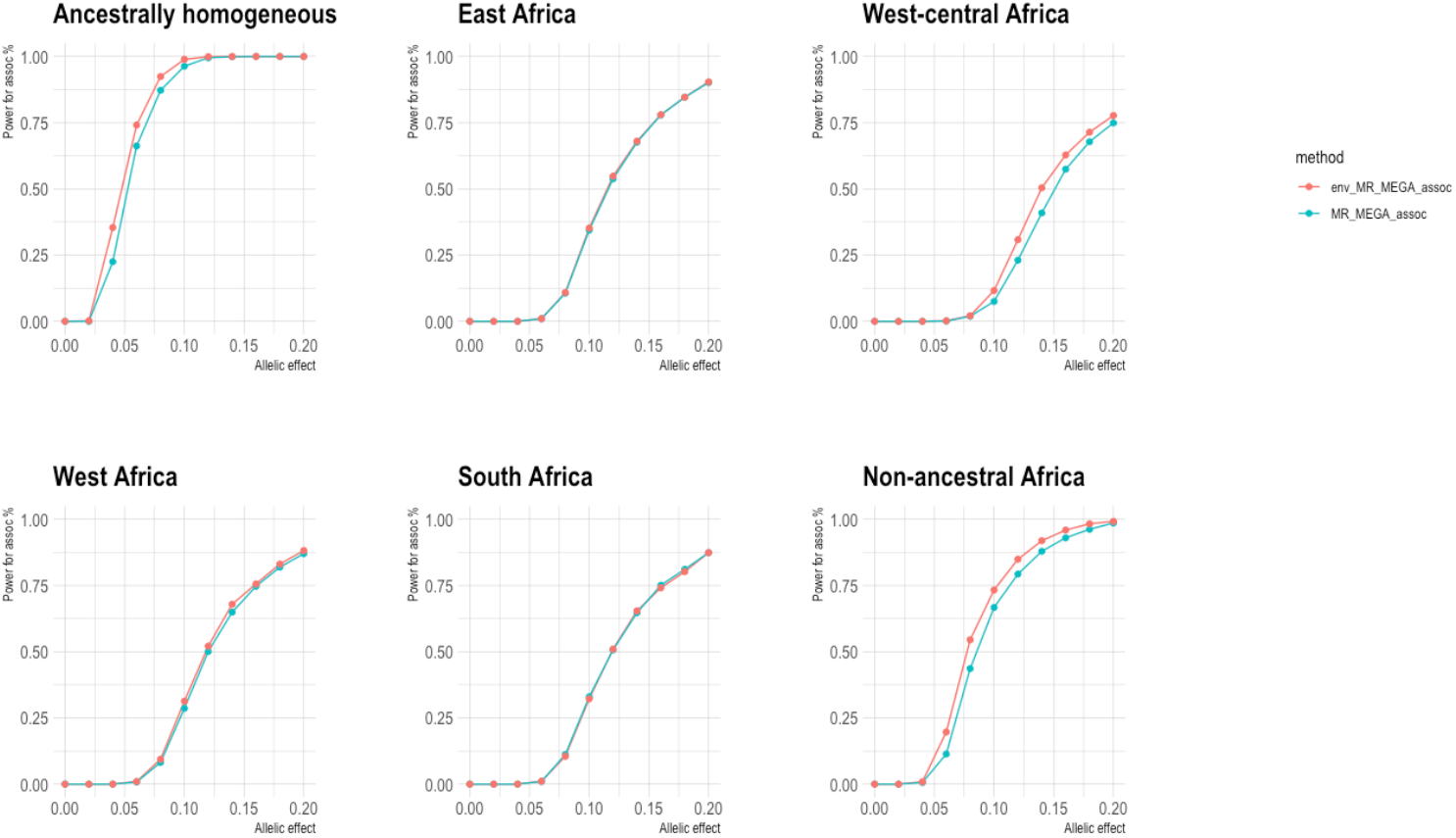
Across six heterogeneity scenarios involving 16 sex-stratified cohorts, where minor random reductions or increases in smoking proportions between male and female cohorts occur (mixed direction), env-MR-MEGA exhibits higher power in detecting association compared to MR-MEGA, especially in ancestrally homogeneous and non-ancestral Africa scenarios. “env_MR_MEGA_assoc” and “MR_MEGA_assoc” refer to the power to detect association obtained from env-MR-MEGA and MR-MEGA. Power was assessed at *P* < *5* × *10*^−*8*^ and based on 1000 simulations with unequal sample sizes.

As the disparity in smoking proportions between males and females increased and the variability in smoking proportions between African regions diminished (**Supplementary Material Table S3**), compared to the first smoking proportion pattern (**Fig2, Supplementary Material Fig S4**, **Fig S5**), the difference in power to detect association between env-MR-MEGA and MR-MEGA in the same direction became more apparent (**Fig 3**). Compared with the same direction setting where all female cohorts have lower smoking proportions, the mixed direction setting includes South Africa and west-central Africa scenarios where one female cohort has a higher smoking proportion. Therefore, in the South Africa and West Central Africa scenarios, the power for association attained from env-MR-MEGA in the same direction setting was notably higher than that in the mixed direction setting. However, compared to the gap of power for association from env-MR-MEGA between same direction and mixed direction settings, the power for association from MR-MEGA varied less between the same direction and mixed direction settings.

**Figure 3.**
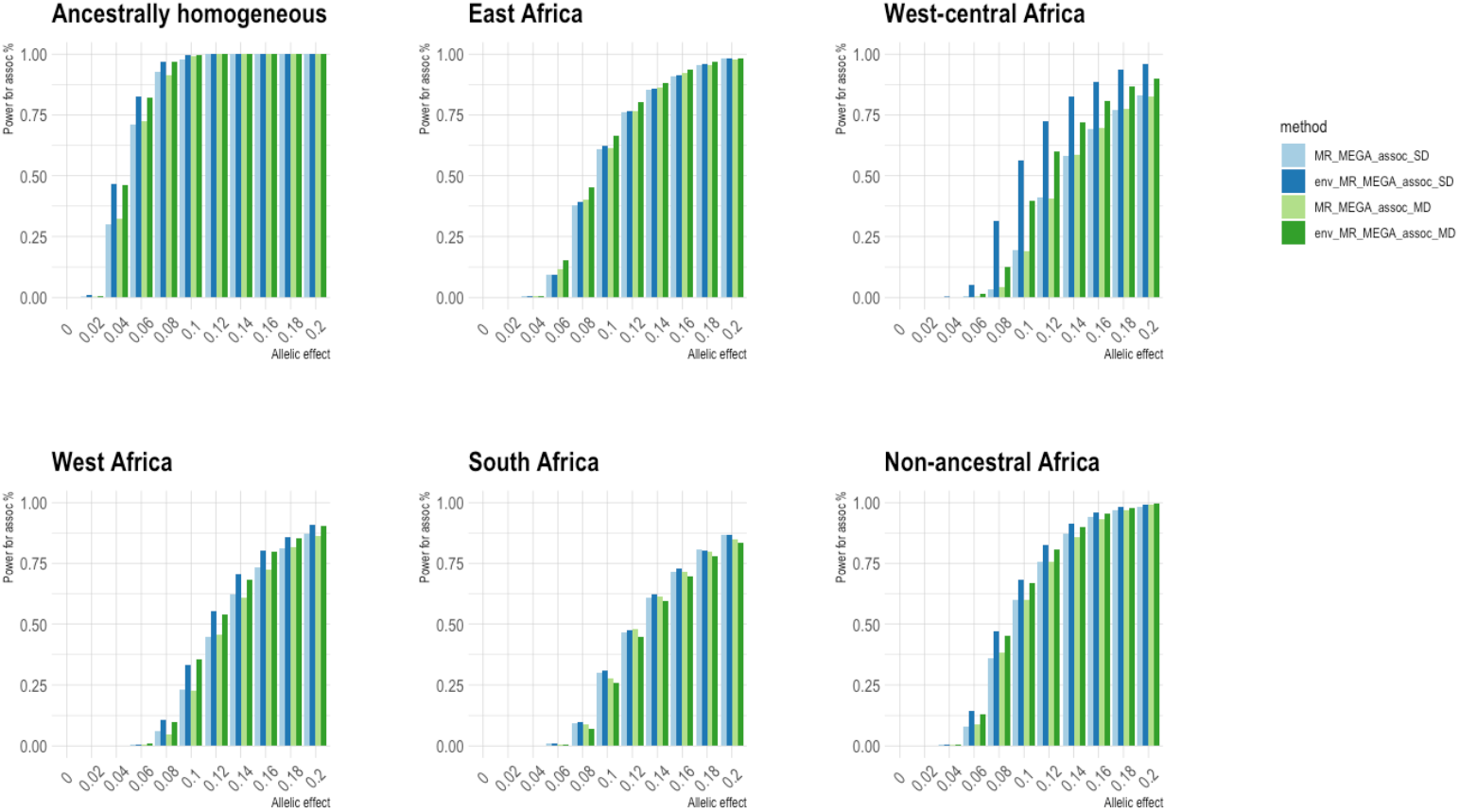
When smoking proportions notably differed between male and female cohorts, env-MR-MEGA gained greater power to detect association compared to MR-MEGA. In the mixed direction pattern, where not all male cohorts share high smoking proportions, both West-central Africa and South Africa scenarios contain one male cohort with a lower smoking proportion. Within West-central Africa and South Africa scenarios, compared the same direction and the mixed direction patterns, the gains in power to detect association by env-MR-MEGA in the same direction pattern exceeded those in the mixed direction pattern. “MR_MEGA_assoc_SD” and “env_MR_MEGA_assoc_SD” correspond to the same direction pattern, while “MR_MEGA_assoc_MD” and “env_MR_MEGA_assoc_MD” correspond to the mixed direction pattern. Power was assessed at *P* < 5 × 10^−8^ and based on 1000 simulations with unequal sample sizes.

In the first pattern, where the smoking proportions varied slightly between male and female cohorts (**Supplementary Material Table S2**), the power to detect heterogeneity due to ancestry and/or environment does not vary dramatically across the three settings for smoking proportions: no-difference, same direction and mixed direction (**Supplementary Material Fig S6**, **Fig S7 and Fig S8**). For the ancestrally homogeneous, west-central Africa and non-ancestral Africa scenarios, where heterogeneity in allelic effect is lower correlated with ancestry compared to smoking exposures, the power for allelic heterogeneity due to ancestry and environment from env-MR-MEGA was notably greater than that due to ancestry from MR-MEGA. For the heterogeneity scenarios where heterogeneity in allelic effects is moderately correlated with ancestry and smoking exposures, east Africa and West Africa, there was not much difference between the power for heterogeneity due to ancestry and the power for heterogeneity due to ancestry and environment. Especially for the South Africa scenario, where the heterogeneity in allelic effect is more strongly correlated with the South Africa ancestry compared to smoking exposures, the power for allelic heterogeneity due to ancestry attained from both env-MR-MEGA and MR-MEGA is noticeably greater than that due to ancestry and environment.

For the scenarios where heterogeneity in allelic effect is correlated with ancestry and smoking exposures (east Africa, west Africa, west-central Africa and South Africa), the power for allelic heterogeneity due to environment from env-MR-MEGA was the lowest amongst all heterogeneity tests in the first smoking pattern (**Supplementary Material Table S2**). As the difference in smoking proportions between female and male cohorts increased and variation in smoking proportions between ancestries diminished (**Supplementary Material Table S3**), the power for heterogeneity due to the environment would increase significantly, particularly for the west-central Africa scenario (**Supplementary Material**, **Fig S9 and Fig S10**). In addition, compared to the mixed direction setting where some female cohorts have higher smoking proportions than male cohorts in the same populations, the same direction in smoking proportions gained more power for heterogeneity in environmental effects than mixed direction.

### Identification of sources of genetic heterogeneity in African cohorts

We applied the env-MR-MEGA method to the summary statistics of twelve sex-stratified African GWAS (representing West Africa, East Africa and South Africa) for LDL-cholesterol using sex, mean BMI, and the proportion of study participants with urban status as environmental variables. We used these variables individually and in pairs as environmental factors. We considered SNPs that have previously attained genome-wide significance for LDL-cholesterol in the meta-analysis of African American GWAS from the GLGC^38^. For these SNPs, we used env-MR-MEGA to investigate the contribution of ancestry and environmental variables to heterogeneity in allelic effects between 12 sex-stratified GWAS from continental Africa. We applied env-MR-MEGA to summary statistics from these studies to assess the impact of sex, BMI, and urban environment on heterogeneity.

The axes of genetic variation show three main clusters, coinciding with East Africa (AWI-Gen East, Uganda), West Africa (AWI-Gen West), and South Africa (AWI-Gen South, Zulu) (**Fig 4a**). Twelve SNPs (**Table 1**) showed allelic heterogeneity (p-value < 0.05) due to ancestry, environment only (sex, BMI, and urban status) and a combination of both ancestry and environmental variables.

**Table 1.**
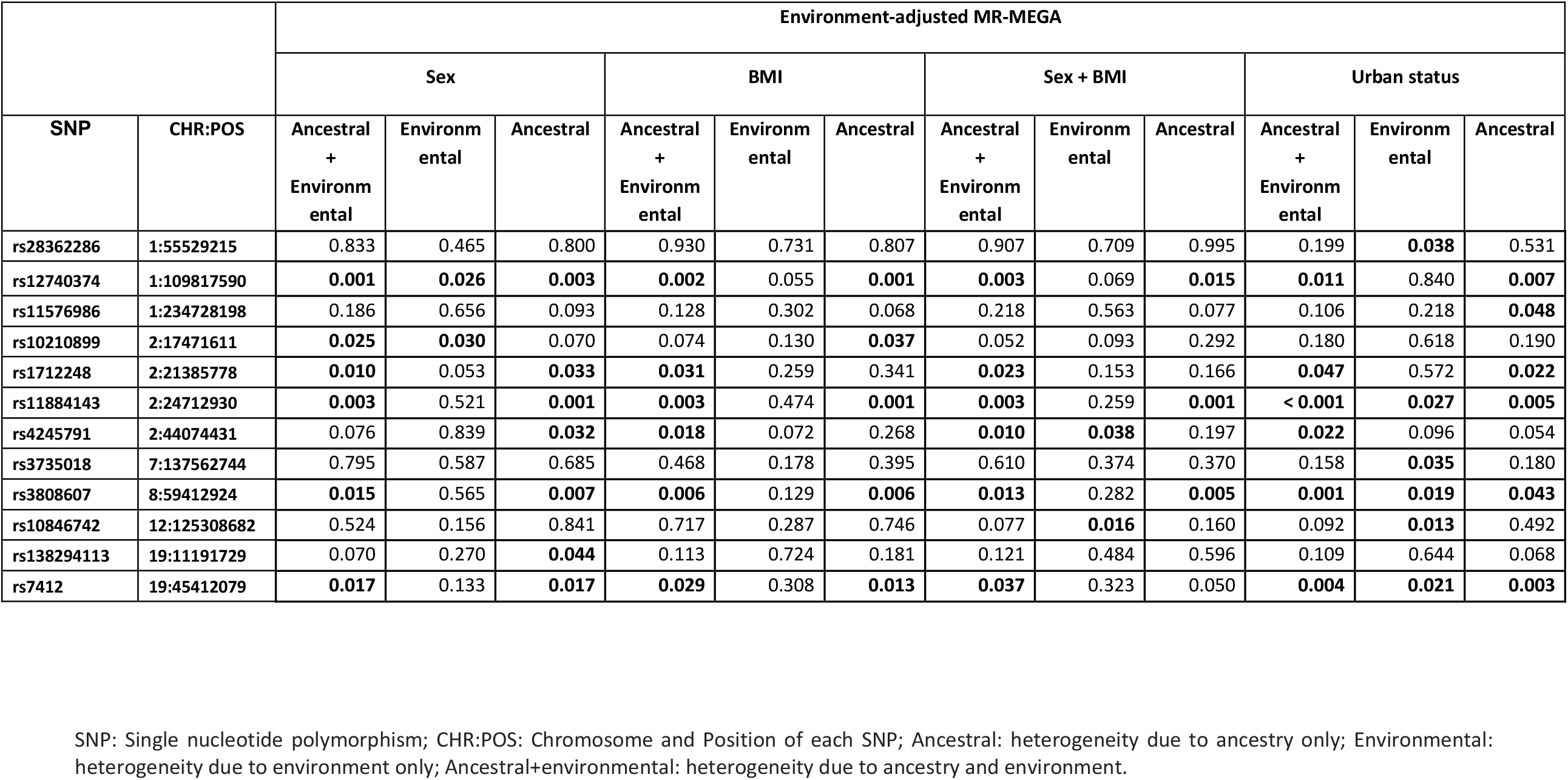
Heterogeneity due to ancestry and environmental variables as identified by 1§env-MR-MEGA. The data presented are the heterogeneity p-values for the env-MR-MEGA (environment-adjusted *Meta*-Regression of Multi-AncEstry Genetic Association) methods. Heterogeneity p-values for sex, BMI (body mass index), sex + BMI, urban status were generated using the env-MR-MEGA methods. The heterogeneity p-values < 0.05 are in bold font.

|  |  | Environment-adjusted MR-MEGA |  |  |  |  |  |  |  |  |  |  |  |
| --- | --- | --- | --- | --- | --- | --- | --- | --- | --- | --- | --- | --- | --- |
|  |  | Sex |  |  | BMI |  |  | Sex + BMI |  |  | Urban status |  |  |
| SNP | CHR:POS | Ancestral + Environm ental | Environm ental | Ancestral | Ancestral + Environm ental | Environm ental | Ancestral | Ancestral + Environm ental | Environm ental | Ancestral | Ancestral + Environm ental | Environm ental | Ancestral |
| rs28362286 | 1:55529215 | 0.833 | 0.465 | 0.800 | 0.930 | 0.731 | 0.807 | 0.907 | 0.709 | 0.995 | 0.199 | **0.038** | 0.531 |
| rs12740374 | 1:109817590 | **0.001** | **0.026** | **0.003** | **0.002** | 0.055 | **0.001** | **0.003** | 0.069 | **0.015** | **0.011** | 0.840 | **0.007** |
| rs11576986 | 1:234728198 | 0.186 | 0.656 | 0.093 | 0.128 | 0.302 | 0.068 | 0.218 | 0.563 | 0.077 | 0.106 | 0.218 | **0.048** |
| rs10210899 | 2:17471611 | **0.025** | **0.030** | 0.070 | 0.074 | 0.130 | **0.037** | 0.052 | 0.093 | 0.292 | 0.180 | 0.618 | 0.190 |
| rs1712248 | 2:21385778 | **0.010** | 0.053 | **0.033** | **0.031** | 0.259 | 0.341 | **0.023** | 0.153 | 0.166 | **0.047** | 0.572 | **0.022** |
| rs11884143 | 2:24712930 | **0.003** | 0.521 | **0.001** | **0.003** | 0.474 | **0.001** | **0.003** | 0.259 | **0.001** | **< 0.001** | **0.027** | **0.005** |
| rs4245791 | 2:44074431 | 0.076 | 0.839 | **0.032** | **0.018** | 0.072 | 0.268 | **0.010** | **0.038** | 0.197 | **0.022** | 0.096 | 0.054 |
| rs3735018 | 7:137562744 | 0.795 | 0.587 | 0.685 | 0.468 | 0.178 | 0.395 | 0.610 | 0.374 | 0.370 | 0.158 | **0.035** | 0.180 |
| rs3808607 | 8:59412924 | **0.015** | 0.565 | **0.007** | **0.006** | 0.129 | **0.006** | **0.013** | 0.282 | **0.005** | **0.001** | **0.019** | **0.043** |
| rs10846742 | 12:125308682 | 0.524 | 0.156 | 0.841 | 0.717 | 0.287 | 0.746 | 0.077 | **0.016** | 0.160 | 0.092 | **0.013** | 0.492 |
| rs138294113 | 19:11191729 | 0.070 | 0.270 | **0.044** | 0.113 | 0.724 | 0.181 | 0.121 | 0.484 | 0.596 | 0.109 | 0.644 | 0.068 |
| rs7412 | 19:45412079 | **0.017** | 0.133 | **0.017** | **0.029** | 0.308 | **0.013** | **0.037** | 0.323 | 0.050 | **0.004** | **0.021** | **0.003** |

SNP: Single nucleotide polymorphism; CHR:POS: Chromosome and Position of each SNP; Ancestral: heterogeneity due to ancestry only; Environmental: heterogeneity due to environment only; Ancestral+environmental: heterogeneity due to ancestry and environment.

**Figure 4.**
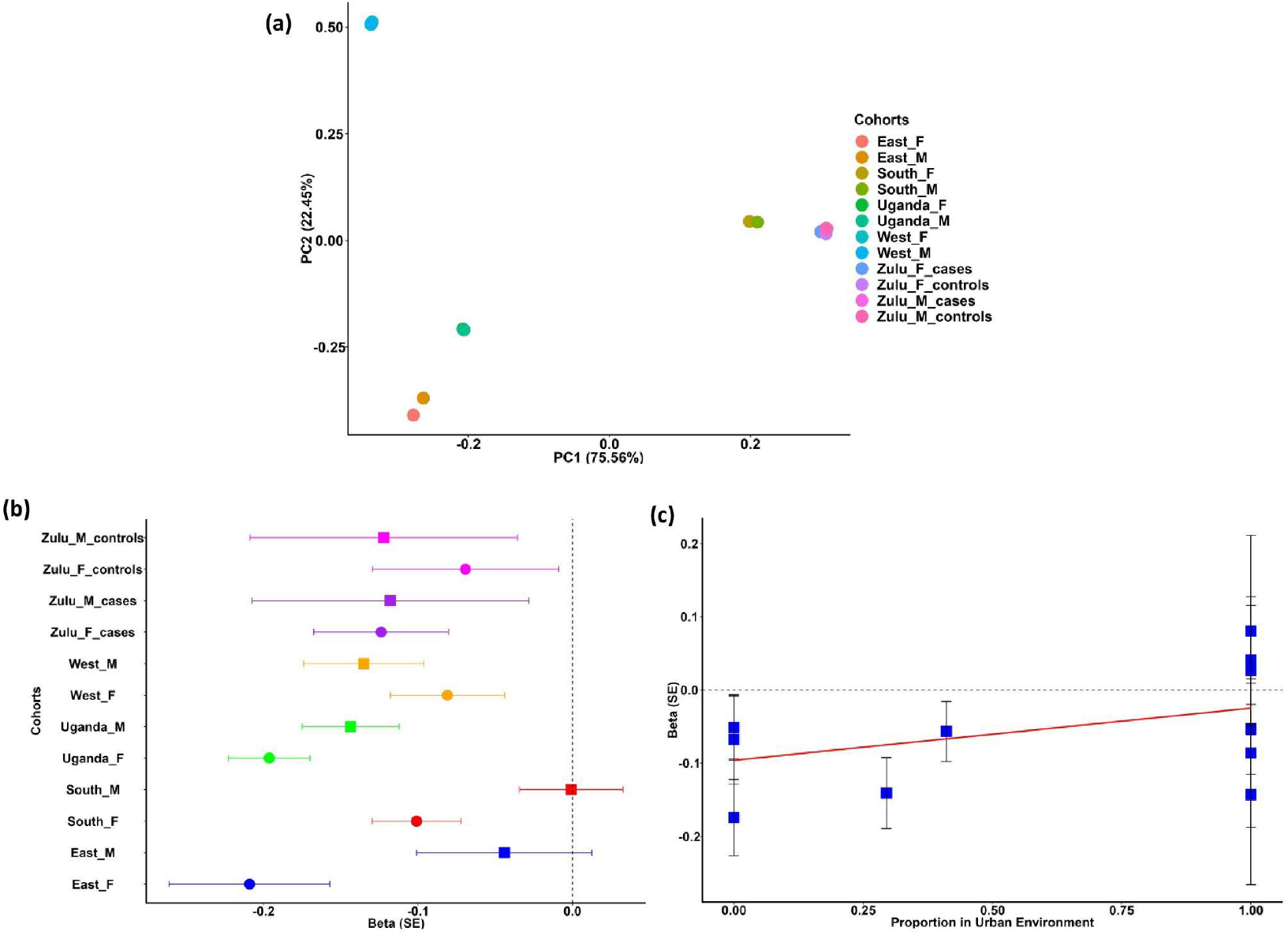
The effect of heterogeneity resulting from ancestry and environmental variables. **(a)** the axes of genetic variation: axes of genetic variation generated based on the allele frequency, showcasing three distinct separations representing cohorts from West Africa (West_M & West_F), East Africa (East_M & East_F and Uganda_M & Uganda_F), and South Africa (South_M & South_F, controls for Zulu_M & Zulu_F and cases for Zulu_M and Zulu_F). **(b)** heterogeneity due to sex with the female cohorts (for Zulu_cases, South, Uganda and East) exhibiting greater effect sizes for rs12740374 than males. **(c)** heterogeneity due to urban status with effect size for rs3735018 increasing towards urban residents’ cohorts. In (b) and (c) 95% confidence intervals are provided for each effect estimate.

Our env-MR-MEGA method detected allelic heterogeneity involving environmental effects in nine SNPs: two SNPs driven by sex (rs12740374 and rs10210899), one SNP jointly driven by combined sex and BMI (rs4245791), five SNPs driven by urban status (rs11884143, rs3735018, rs3808607, rs28362286 and rs7412), and one SNP driven jointly by sex and BMI, as well as urban status (rs10846742); rs28362286 and rs7412 were genome-wide significant SNPs for LDL in twelve African cohorts overlapping with the GLGC meta-analysis^38^ (**Supplementary Data 1**). Female cohorts from East Africa are most strongly associated with decreased LDL-cholesterol (**Figure 4b**), while those residing in urban areas are most strongly associated with increased LDL-cholesterol (**Figure 4c**). Three SNPs displayed heterogeneity due to ancestry alone (rs11576986,rs1712248 and rs138294113) and a further four SNPs had heterogeneity due to both ancestry and an environmental effect (rs12740374, rs3808607, rs11884143 and rs7412) (**Supplementary Data 1**).

## Discussion

We have developed an environment-adjusted meta-regression of GWAS, env-MR-MEGA, utilising GWAS summary statistics, along with the environmental exposure and ancestry. The model is built upon the framework of MR-MEGA^30^, treating the allelic effects as a function of environmental exposures in addition to genetic ancestry. Through a series of simulations, we demonstrate that env-MR-MEGA outperforms MR-MEGA in terms of power to detect association when there is indeed an effect of the environmental exposure on heterogeneity, as well as the power to detect heterogeneity that is due to various sources, i.e. genetic ancestry and/or environmental factors.

The framework of env-MR-MEGA not only accounts for ancestry obtained by deriving axes of genetic variation via multi-dimensional scaling but also study-level environmental impacts obtained by taking the mean or proportion of the individual-level environmental data within each cohort. Our simulation studies and African data application focused on cohorts from different African regions, which include East Africa, West Africa, and South Africa. In our analyses, we have focused on the two axes of genetic variation and illustrated that the two axes of genetic variation are sufficient to distinguish these cohorts within the African continent (**Supplementary material Fig S1, Fig 4a**). The environmental exposures were given as the sex-stratification variable, the mean BMI within each cohort, and the study-level urban status proportions. By incorporating genetic variation axes and study-level environmental exposures as covariates in the meta-regression model, env-MR-MEGA can detect heterogeneity in allelic effects attributed to ancestry and environmental factors, respectively.

When the sex-stratification variable is treated as the environmental exposure only, our simulation shows that if allelic effects are correlated with sex and ancestry, env-MR-MEGA gains the greatest power to detect association over MR-MEGA across all scenarios, as expected. Additionally, as the correlation between allelic heterogeneity and ancestry decreases, such as in the ancestrally homogeneous scenario and the non-ancestral scenario, the test for allelic heterogeneity due to environmental effect gains more power than that due to ancestry (**Supplementary Material**, **Figure S2**). Applying env-MR-MEGA to association studies of LDL-cholesterol highlights notable evidence of heterogeneity in sex, which was observed at rs12740374 and rs10210899 (**Supplementary Data 1**).

Our simulation results demonstrate that when the heterogeneity in allelic effects is lower correlated with ancestry compared to the environmental exposures, such as ancestrally homogeneous, west-central Africa and non-ancestral Africa scenarios, env-MR-MEGA gains noticeably higher power to detect association over MR-MEGA (**Fig 2, Supplementary Material Fig S4**, **Fig S5**). This improvement in power is attributed to the weak correlation between allelic heterogeneity and ancestry. Specifically, as the correlation between allelic heterogeneity and ancestry weakens, the power for heterogeneity due to ancestry and environment from env-MR-MEGA became notably greater than that due to ancestry alone from MR-MEGA.

In the application to GWAS meta-analysis of LDL-cholesterol across African GWAS, BMI and urban status are treated as environmental exposures in addition to sex-stratification. Our assessment of heterogeneity of genome-wide significant SNPs in the GLGC study revealed nine SNPs with environmental heterogeneity of which (four SNPs were driven by both ancestry and environmental variables. Three SNPs displayed heterogeneity due to ancestry only.

Compared to MR-MEGA, the key advantage of env-MR-MEGA is that the environment-adjusted meta-regression model accounts for heterogeneity in allelic effects that can be attributed to ancestry and environmental factors instead of ancestry alone. When there is heterogeneity in allelic effects that is not correlated with ancestry, we can quantify the contribution of environmental effects to the heterogeneity of allelic effects between GWAS. Consequently, env-MR-MEGA offers a more comprehensive approach to the discovery across GWAS from diverse populations. With the increasing availability of GWAS from more diverse populations, an efficient statistical methodology that will account for heterogeneity in meta-analysis of GWAS, such as env-MR-MEGA, shows great promise for future improvements in our understanding of the genetic basis of complex human traits.

## Acknowledgements

This research is funded by the UK Medical Research Council (MR/W02098X/1). SW and JA are also funded by the UK Medical Research Council (MC_UU_00002/4). For the purpose of Open Access, the authors have applied a CC BY public copyright licence to any Author Accepted Manuscript version arising from this submission

## Materials and Methods

Following MR-MEGA^30^, consider the *K* sets of GWAS summary statistics of a complex trait from *K* cohorts and assume that allelic effects for all cohorts are aligned to the same reference allele at each variant. For the *k*th study, we denote by *p*_*kj*_ the reference allele frequency of the *j*th SNP in the *k*th cohort. We derive a matrix of pairwise Euclidean distances between cohorts from a subset of autosomal variants, denoted *D* = [*d*_*kk*_′]. More specifically, each component *d*_*kk*_′ of the matrix *D* is given by 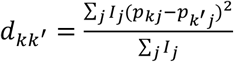, where *I*_*j*_ is a binary variable indicating whether the *j*th variant is in the subset of SNPs used for calculating the distance matrix.

In the procedure of generating the subset to derive the distance matrix, we follow the strategy of MR-MEGA^30^. First, the autosomal variants from the reference panel with minor allele frequency (MAF) >5% in all cohorts are retained and are divided into 1Mb bins. Then we randomly select one variant from each bin and aggregate them to calculate the pairwise Euclidean distances.

We denote *T* axes of genetic variation across *K* cohorts by *x*_*t*_ = (*x*_1*t*_, *x*_2*t*_, …, *x*_*kt*_), *t* = 1, …, *T*, which are derived from the multi-dimensional scaling of the distance matrix. Taking account that the environmental exposures could be specific to each cohort, we construct the cohort-level environmental covariates by taking the mean or proportion of the individual-level environmental data within each cohort. Here we denote *S* cohort-level environmental covariates across K GWAS by *z*_*s*_ = (*z*_1*s*_, *z*_2*s*_, …, *z*_*ks*_), *s* = 1, …, *S*. It is noted that the number of axes of genetic variation and the number of environmental covariates would depend on the number of cohorts, but the limitation of the number of axes of genetic variation and the number of environmental covariates should be *T* + *S* < *K* − 2.

Following the notation for MR-MEGA, we denote the estimated allelic effect and the corresponding variance of the *k*th cohort at the *j*th variant by *b*_*kj*_ and *ν*_*kj*_. For the *j*th variant, our environment-adjusted meta regression model is given by

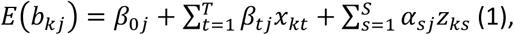

where *β*_*tj*_ represents the effect of the *t*th axis of genetic variation for the *j*th variant and α_*sj*_ represents the effect of the *s*th environmental covariate for the *j*th variant. In the model, *β*_0*j*_ is the intercept for the *j*th variant and can be interpreted as the expected allelic effect of variant *j* when all axes of genetic variation and environmental covariates are zero.

To test the null hypothesis of no association of the *j*th variant across all cohorts, we compare the null model with *β*_0*j*_ = *β*_1*j*_ = ⋯ = *β*_*Tj*_ = α_1*j*_ = ⋯ = α_*sj*_ = 0 in model (1) to that without constrained parameters. The resulting test statistic for the *j*th variant has chi-square distribution with (*T* + *S* + 1) degrees of freedom. To test the presence of heterogeneity due to ancestry and environment effect, we would compare the model with *β*_1*j*_ = ⋯ = *β*_*Tj*_ = α_1*j*_ = ⋯ = α_*sj*_ = 0 in model (1) to that without constrained parameters. The corresponding test statistic has an approximate chi-square distribution with (*T* + *S*) degrees of freedom. We also test the heterogeneity due to ancestry and environment separately. For the heterogeneity in allelic effects, the test statistic would be obtained by comparing the model with *β*_1*j*_ = ⋯ = *β* _*Tj*_ = 0 to that without constrained parameters, which follows an approximate chi-square distribution with *T* degrees of freedom. In the same way, we can test for heterogeneity in environmental effects by comparing the model with α_1*j*_ = ⋯ = α_*sj*_ = 0 and the model without constrained parameters, with the test statistic following the approximate chi-square distribution with *S* degrees of freedom. Finally, after accounting for ancestry and environment, we can test residual heterogeneity in allelic effects by the deviance of the model (1) with unconstrained parameters. The corresponding resulting test statistic has an approximate chi-square distribution with (*K* − *S* − *T* − 1) degrees of freedom.

## Simulation study design

We conducted extensive simulation studies to compare the performance of env-MR-MEGA and MR-MEGA in terms of statistical power to detect genetic association and heterogeneity due to environment and ancestry across genetically diverse populations from Africa. Here, assuming Hardy-Weinberg equilibrium, we simulated genotypes from the allele frequencies of eight populations from Africa: two Zulu cohorts from southern Africa, Durban Diabetes study (DDS) and Durban Case Control study (DCC)^35^, the Uganda Genome Resource^36^ and Esan (ESN), Gambian (GWD), Luhya (LWK), Mende (MSL), and Yoruba (YRI) from Phase 3 of 1000 Genomes^37^ (**Supplementary Material Table S1)**.

We considered two general settings in our simulations of 16 cohorts, sex-stratified across the eight populations from Africa: (i) heterogeneity due to genetic ancestry and sex (different model for each sex); (ii) heterogeneity due to genetic ancestry and smoking status (same model for both sexes).

In our simulations where allelic effects are correlated with ancestry and sex, we set: *Y* = *ε* for males and *Y* = *β* × *g* + *ε* for females, where *g* denotes the genotype for one genetic variant which would be coded 0, 1 or 2 representing the number of copies of the minor allele, *β* is the effect size for the genetic variant and *ε*∼*N*(0, *sd*), *sd* = 0.45. Here, we treat sex as an environmental covariate that acts as an indicator variable since we stratify population groups by sex. In our simulations where there is no sex effect and heterogeneity in allelic effects is correlated with ancestry and smoking status, we set: *Y* = *ε* for non-smokers; *Y* = *β* × *g* + *ε* for smokers, where *ε*∼*N*(0, *sd*), *sd* = 0.45.

We considered six heterogeneity scenarios across population groups, parameterised in terms of allelic effects *β* in each population group (**Supplementary Material Table S1**). These heterogeneity scenarios could be summarised into three groups: ancestrally homogeneous, ancestry-specific, and non-ancestral. For the ancestrally homogeneous setting, the allelic effect is the same across all African sub-populations (i.e. no ancestral heterogeneity). In the population-specific setting, the allelic effect is specific to sub-populations in the specified geographic region (east Africa, west-central Africa, west Africa, or south Africa). Finally, for the non-ancestral setting, the allelic effect is specific to one sub-population in each African region, i.e. heterogeneity within each African region. Details of specific simulation studies follow.

### Simulation study 1: heterogeneity in allelic effect is correlated with sex and population groups

Considering an environment-adjusted meta-analysis of eight GWAS of a quantitative trait, we initially assumed that the eight sub-populations from Africa, ESN, YRI, GWD, MSL, DCC, DDS, LWK and UGD, have unequal sample sizes of 8000, 6500, 7000, 8000, 6000, 6000, 6500 and 8000, respectively. We also stratified each sub-population by sex. For simplicity, we assumed equal sample sizes for male and female cohorts from the same sub-population. We considered a model of female-specific effect (*β* = 0 in male cohorts) for the above-mentioned six heterogeneity scenarios (**Supplementary Material Table S1**). Additionally, we specifically considered the homogeneity in sex and ancestry scenario, where male and female cohorts share the same model *Y* = *β* × *g* + *ε* across all cohorts, to assess the performance of env-MR-MEGA in terms of type I error. In env-MR-MEGA, the environment covariate was constructed by a binary vector, each component consisting of 1 for female and 0 for male.

To evaluate the power and type I errors for each heterogeneity scenario, we generated 1000 replications. Within each simulation replication, we randomly selected a causal variant with MAF>1% under Hardy-Weinberg equilibrium (HWE) assumption in all populations using the population-specific allele frequency for the population from the 1000 Genomes Project. As a baseline power comparison between the two methods, we considered equal-sized cohorts (1000 in each female/male cohort) and then unequal sample sizes.

### Simulation study 2: the heterogeneity in allelic effect is correlated with the environmental covariate and population groups

To investigate the impact of the environmental effect, we considered the setting in which the heterogeneity in allelic effect is correlated with the environmental factor. Here, smoking status was treated as the environmental covariate, and the proportion of smokers varied across the 8 sub-populations, including the sex-stratified 16 male/female cohorts. Here, we replicated the procedure outlined in the sex-stratified simulations to simulate genotype data. The sample sizes and 1000 causal variants were consistent with those in the sex-stratified simulations for this simulation. Under the setting of the heterogeneity of allelic effects correlated with the environmental effect, we examined a model of smoker-specific association with the trait; individuals who are smokers within each cohort have a genetic effect, while those who do not smoke have no genetic effect. The environmental covariate is a vector of smoking proportions for the corresponding female and male cohorts in the associated environment-adjusted meta-regression model.

We considered three different settings for the difference in smoking proportions between female and male cohorts: (a) no-difference: female and male cohorts in the same population shared the same smoking rate; (b) same direction: smoking rates in female cohorts were always lower than that in male cohorts; (c) mixed direction: among the 8 sub-populations, not all female smoking rates were lower than that in male cohorts. To make these settings more general and to cover a range of scenarios, we constructed two patterns for smoking proportions: (i) slight differences in smoking proportions between males and females but large differences in smoking proportions between populations in the same African region; (ii) large differences in smoking proportions between male and females but minor differences in smoking proportions between populations in the same African region (**Supplementary Material Table S2 and Table S3**). For example, the proportions of smokers vary greatly between females and males^39^, whereas other covariates may not differ as much between females and males.

### Simulations: Power Evaluation

Within the two simulation studies, we tested for genetic association using env-MR-MEGA, with two axes of genetic variation and sex and environmental covariates. For comparison, we also tested for association using MR-MEGA with two axes of genetic variation. The power to detect association was assessed at a genome-wide significance level (*P* < 5 × 10^−8^). For each environment-adjusted meta-regression model, we also obtained the power for detecting allelic heterogeneity due to ancestry and environment, as well as the power for allelic heterogeneity due to environment and the power for allelic heterogeneity due to ancestry alone. For each meta-regression model, we obtained the power to detect ancestry-correlated heterogeneity. In all replicates, power for heterogeneity was evaluated at nominal significance (*P* < 0.05).

### Application of env-adjusted MR-MEGA

To demonstrate the efficacy of a novel method for assessing and controlling heterogeneity, we employed the env-MR-MEGA approach on genome-wide association study (GWAS) summary statistics for LDL-cholesterol. The GWAS data was stratified by sex and assembled from diverse sub-Saharan African cohorts, comprising a total of 19,589 participants (in AWI-Gen cohort, South African Zulu cohorts, and Uganda Genome Resource).

### AWI-Gen cohort

This study utilised 10,898 participants from the Africa Wits-INDEPTH partnership for Genomics studies (AWI-Gen), who passed the post-imputation filters. The AWI-Gen cohort comprises six centres in sub-Saharan African countries, including three centres in South Africa (denoted as South Africa in this study: South male = 2225, South female = 3040), one centre from Kenya (referred to as East Africa: East male = 805, East female = 961), and two centres from West Africa (one from Ghana and one from Burkina Faso, denoted as West Africa: West male = 1883, West female = 1984). EAGLE2 was used for pre-phasing, and the default Positional Burrows–Wheeler Transform (PBWT) algorithm was used for imputation. Imputation was performed on the cleaned dataset with 1,729,661 SNPs remaining after quality control, which included removing closely related individuals) using the African Genome Resources reference panel in the Sanger Imputation Server. After imputation, poorly imputed SNPs with info scores less than 0.6, SNPs with minor allele frequency (MAF) < 0.01, and Hardy-Weinberg Equilibrium (HWE) p-value < 0.00001 were excluded. The final quality-controlled imputed data had 13.98 M SNPs. Further information on AWI-Gen cohorts has been reported in previous studies^40, 41^. The study analysed the fasting serum lipids of the participants using a colorimetric assay (Randox Plus clinical chemistry analyser: Crumlin, Northern Ireland, UK). Low-density lipoprotein cholesterol (LDL-C) was then calculated using the Friedewald equation^42^ and measured in mmol/L. LDL-cholesterol was regressed on age, and the first five principal components and the residuals were transformed by inverse rank normalisation. Using the residuals stratified by sex and centre, a linear mixed model association analysis was implemented in Genome-wide Complex Trait Analysis (GCTA version 1.91.7 beta1)^43^.

### South African Zulu cohorts

The cohort of South African Zulu comprises 2707 individuals from the Durban Diabetes Study (DDS) and the Durban Diabetes Case Control Study (DCC). Previous studies have provided details on the participants’ demographics, genotyping platforms, and imputation procedures^44^. After quality control, we used 16,559,897 variants and 2572 individuals. 1599 participants were cases (Zulu male cases = 375; Zulu female cases = 1224) and 973 controls (Zulu male controls = 298; Zulu female controls = 675) for the association analysis. Fasting LDL-cholesterol levels (mmol/L) were measured using the ABBOTT ARCHITECT 2: CI 8200 automated analyser, which employs an enzymatic method. To conduct the association testing, we stratified the cases and controls separately by sex and regressed LDL-cholesterol levels on age, and the first two principal components; the residuals were transformed by inverse rank normalisation, and association analysis was implemented in Genome-wide Complex Trait Analysis software (GCTA version 1.91.7 beta1)^43^.

### Uganda Genome Resource

The Uganda Genome Resource dataset, consisting of 7,000 individuals, has been documented in recent years^45,46^. This documentation covers the genotyping platforms, quality control measures, and imputation techniques. The final dataset used for genome-wide association testing, which excluded MAF < 1%, contained 15,783,409 SNPs for 6,119 individuals (Uganda male = 2618; Uganda female = 3501). Non-fasting low-density lipoprotein (LDL)-cholesterol levels (mmol/L) were measured directly using the colorimetric assay described by Sugiuchi et al^47^. LDL-cholesterol was regressed on age, and the first two principal components and the residuals were transformed by inverse rank normalisation. A sex-stratified association analysis was conducted on the residuals using a mixed linear model in Genome-wide Complex Trait Analysis (GCTA version 1.91.7 beta1)^43^.

### Environment-adjusted MR-MEGA method

To generate axes of genetic variation, we applied a filter to exclude SNPs with MAF below 5%. Our env-adjusted MR-MEGA (env-MR-MEGA) method incorporated three variables to account for potential environmental factors that could introduce heterogeneity: sex, BMI, and participants’ settings (urban or rural). As we had cohorts from different settings, we also included the proportion of participants (stratified by sex) residing in urban areas to account for potential heterogeneity in allelic effects. Our sex-stratified meta-analysis involved 12 cohorts and utilised env-MR-MEGA method (for sex, BMI, sex + BMI, urban setting). The output file included p-values for association with LDL-cholesterol, as well as heterogeneity p-values due to ancestry and/or environment.

Within our analyses, we focus on genetic variants that reached genome-wide significance within the admixed African or African genetic ancestry group defined within the Global Lipids Genetic Consortium (GLCG) analyses^38^. The GLGC analysis consisted of 90,000 individuals, which is considerably larger than our analyses, so we treat their identified genome-wide significant variants in the African-American admixed individuals as a “gold standard” and assess allelic heterogeneity at these variants within our cohorts.

## Data availability

1KG cohorts used in the paper are available on MAGMA (https://ctg.cncr.nl/software/MAGMA/ref_data/). UGR and Zulu datasets are available on NHGR-EBI GWAS catalog link (https://www.ebi.ac.uk/gwas/publications/31675503#study_panel). GLGC summary statistics for the lipid traits are available at https://doi.org/10.1038/s41586-021-04064-3 (Supplementary Table 3). The summary statistics from our env-MR-MEGA analyses of twelve sex-stratified GWAS from Africa, at the genome-wide significant variants for LDL in the African-American meta-analysis from GLGC, are available in Supplementary Material, Supplementary Data 1.

## Code Availability

Our proposed env-MR-MEGA method, env-MR-MEGA is freely available as an R library at https://github.com/SiruRooney/env.MRmega (https://zenodo.org/records/11047160).

