## Supplementary Material for "Accounting for heterogeneity due to environmental sources in meta-analysis of genome-wide association studies"

**Table S1. Heterogeneity scenarios parameterised in terms of $\beta$ in each reference sub-population. Among eight populations, west-central Africa (ESN,YRI), west Africa (GWD,MSL), one population of east Africa (LWK) were collected from Phase 3 of the 1000 Genomes project; southern Africa (DCC, DDS) and another population of east Africa (UGD) were collected from Uganda Genome Resource.**

| Population | | Heterogeneity scenarios (allelic effect $\beta$) | | | | | |
|---|---|---|---|---|---|---|---|
| Code | Region | Ancestrally Homogeneous | East Africa | West-central Africa | West Africa | South Africa | Non-ancestral Africa |
| ESN | West-central Africa | $\beta$ | 0 | $\beta$ | 0 | 0 | $\beta$ |
| YRI | West-central Africa | $\beta$ | 0 | $\beta$ | 0 | 0 | 0 |
| GWD | West Africa | $\beta$ | 0 | 0 | $\beta$ | 0 | $\beta$ |
| MSL | West Africa | $\beta$ | 0 | 0 | $\beta$ | 0 | 0 |
| DCC | South Africa | $\beta$ | 0 | 0 | 0 | $\beta$ | $\beta$ |
| DDS | South Africa | $\beta$ | 0 | 0 | 0 | $\beta$ | 0 |
| LWK | East Africa | $\beta$ | $\beta$ | 0 | 0 | 0 | $\beta$ |
| UGD | East Africa | $\beta$ | $\beta$ | 0 | 0 | 0 | 0 |

**Table S2. Small difference in smoking proportions between female and male cohorts in which eight populations from different Africa regions were stratified by sex.**

| Populations | | Smoke proportions (female/male) | | |
|---|---|---|---|---|
| Code | Region | Non-difference | Same direction | Mixed direction |
| DCC | South Africa | 0.149/0.149 | 0.088/0.226 | 0.088/0.226 |
| DDS | South Africa | 0.644/0.644 | 0.617/0.719 | 0.719/0.617 |
| UGD | East Africa | 0.659/0.659 | 0.593/0.691 | 0.691/0.593 |
| LWK | East Africa | 0.135/0.135 | 0.12/0.231 | 0.12/0.231 |
| ESN | West-central Africa | 0.655/0.655 | 0.604/0.714 | 0.714/0.604 |
| YRI | West-central Africa | 0.163/0.163 | 0.096/0.202 | 0.096/0.202 |
| GWD | West Africa | 0.676/0.676 | 0.595/0.749 | 0.749/0.595 |
| MSL | West Africa | 0.15/0.15 | 0.075/0.235 | 0.075/0.235 |

**Table S3. Large difference in smoking proportions between female and male cohorts in which eight populations from different African regions were stratified by sex.**

| Populations | | Smoke proportions (female/male) | |
|---|---|---|---|
| Code | Region | Same direction ( lower proportion in female) | Mixed direction |
| DCC | South Africa | 0.128/0.631 | 0.631/0.128 |
| DDS | South Africa | 0.152/0.652 | 0.152/0.652 |
| UGD | East Africa | 0.165/0.876 | 0.876/0.165 |
| LWK | East Africa | 0.243/0.81 | 0.81/0.243 |
| ESN | West-central Africa | 0.215/0.836 | 0.215/0.836 |
| YRI | West-central Africa | 0.061/0.888 | 0.888/0.061 |
| GWD | West Africa | 0.192/0.676 | 0.192/0.676 |
| MSL | West Africa | 0.066/0.655 | 0.066/0.655 |

**Table S4. False positive error rates (Type I errors), at a genome-wide significance threshold ($P < 5 \times 10^{-8}$), to detect association from env-MR-MEGA and MR-MEGA across a range of heterogeneity scenarios are well-calibrated.**

| Heterogeneity scenario | Unequal sample size | | Equal sample size | |
|---|---|---|---|---|
| | env-MR-MEGA | MR-MEGA | env-MR-MEGA | MR-MEGA |
| Ancestrally homogeneous | 0.041 | 0.046 | 0.038 | 0.031 |
| East Africa | 0.041 | 0.046 | 0.038 | 0.031 |
| West-central Africa | 0.041 | 0.046 | 0.038 | 0.031 |
| West Africa | 0.041 | 0.046 | 0.038 | 0.031 |
| South Africa | 0.041 | 0.046 | 0.038 | 0.031 |
| Non-ancestral Africa | 0.041 | 0.046 | 0.038 | 0.031 |

**Table S5**. **In the homogeneity in sex and ancestry scenario, where male and female cohorts share the same allelic effects, tests for heterogeneity in allelic effects due to environment and ancestry from env-MR-MEGA, as well as heterogeneity due to environment and ancestry separately are calibrated as well.** Power to detect heterogeneity due to ancestry and/or environment were assessed at $P < 0.05$. "AEhet" refers to the power to detect heterogeneity due to ancestry and environment, "Ahet" refers to the power to detect heterogeneity due to ancestry alone and "Ehet" refers to the power to detect heterogeneity due to environment alone.

| env-MR-MEGA: unequal sample size | | | |
|---|---|---|---|
| $\beta$ | AEhet | Ahet | Ehet |
| 0 | 0.048 | 0.054 | 0.045 |
| 0.02 | 0.048 | 0.054 | 0.045 |
| 0.04 | 0.048 | 0.054 | 0.045 |
| 0.06 | 0.048 | 0.054 | 0.045 |
| 0.08 | 0.048 | 0.054 | 0.045 |
| 1.00 | 0.048 | 0.054 | 0.045 |
| env-MR-MEGA: equal sample size | | | |
| 0 | 0.049 | 0.040 | 0.053 |
| 0.02 | 0.049 | 0.040 | 0.053 |
| 0.04 | 0.049 | 0.040 | 0.053 |
| 0.06 | 0.049 | 0.040 | 0.053 |
| 0.08 | 0.049 | 0.040 | 0.053 |
| 1.00 | 0.049 | 0.040 | 0.053 |

**Figure S1. Axes of genetic variation separating eight populations from African ancestry.** The first two axes of genetic variation from multi-dimensional scaling of the Euclidean distance matrix between 8 populations are sufficient to separate population groups from different regions of Africa: east Africa (LWK, UGD), south Africa (DCC,DDS), west-central Africa (ESN, YRI) and west Africa (GWD, MSL).

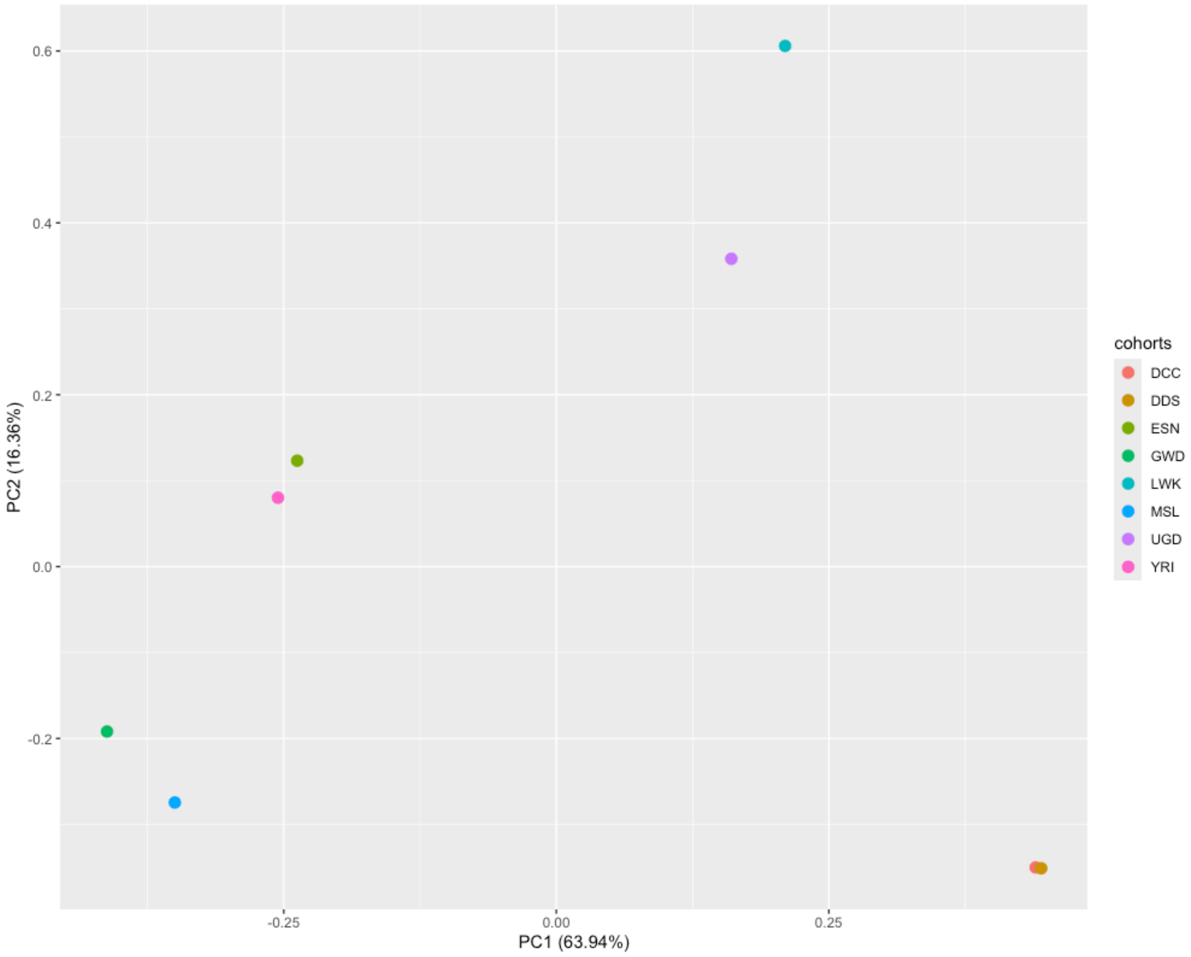

**Figure S2. Across six heterogeneity scenarios, where the 16 sex-stratified cohorts share unequal sample sizes (≥3000 in each female/male cohort), power for heterogeneity of allelic effects due to ancestry and environment from env-MR-MEGA was greater than power for heterogeneity of allelic effects due to ancestry alone from MR-MEGA.** "env_MR_MEGA_AEhet" (red line) corresponds to the power to detect heterogeneity due to ancestry and environment attained from env-MR-MEGA; "env_MR_MEGA_Ahet" (green line) corresponds to the power to detect heterogeneity due to ancestry alone attained from env-MR-MEGA; "env_MR_MEGA_Ehet" (blue line) corresponds to the power to detect heterogeneity due to environment alone attained from env-MR-MEGA; "MR_MEGA_Ehet" (purple line) corresponds to the power to detect heterogeneity due to ancestry obtained from MR-MEGA. Power to detect heterogeneity due to ancestry and/or environment was assessed at $P < 0.05$.

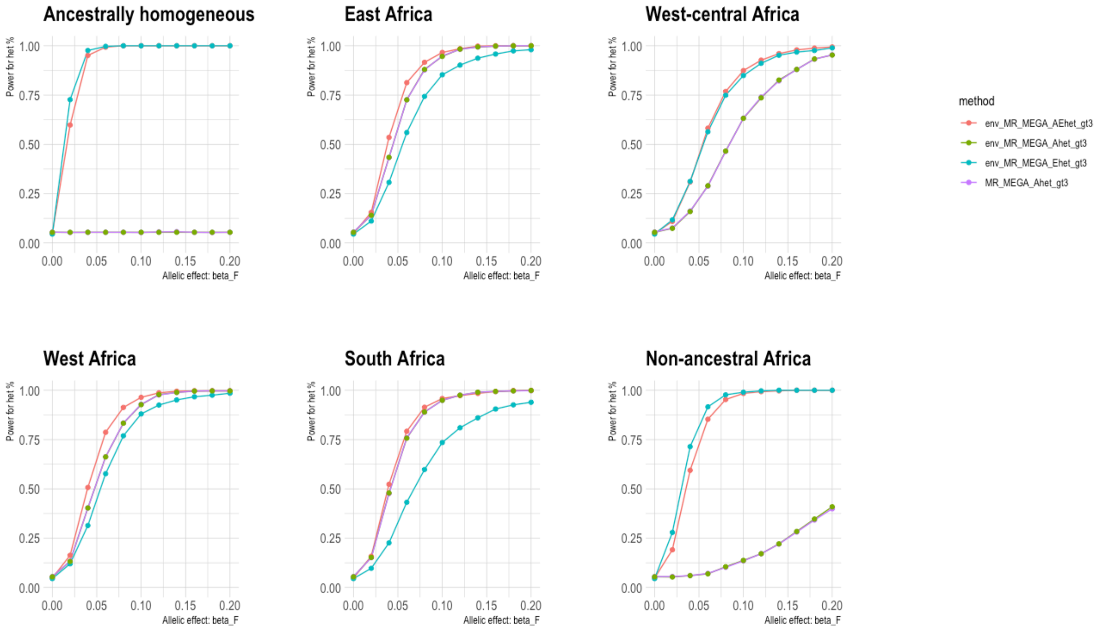

**Figure S3. Across six heterogeneity scenarios where the 16 sex-stratified cohorts share equal sample sizes (1000 in each female/male cohort), power for heterogeneity of allelic effects due to ancestry and environment from env-MR-MEGA was greater than power for heterogeneity of allelic effects due to ancestry and environment from MR-MEGA.** "env_MR_MEGA_AEhet" (red line) corresponds to the power to detect heterogeneity due to ancestry and environment attained from env-MR-MEGA; "env_MR_MEGA_Ahet" (green line) corresponds to the power to detect heterogeneity due to ancestry alone only attained from env-MR-MEGA; "env_MR_MEGA_Ehet" (blue line) corresponds to the power to detect heterogeneity due to environment alone attained from env-MR-MEGA; "MR_MEGA_Ehet" (purple line) corresponds to the power to detect heterogeneity due to ancestry attained from MR-MEGA. Power to detect heterogeneity due to ancestry and/or environment was assessed at $P < 0.05$.

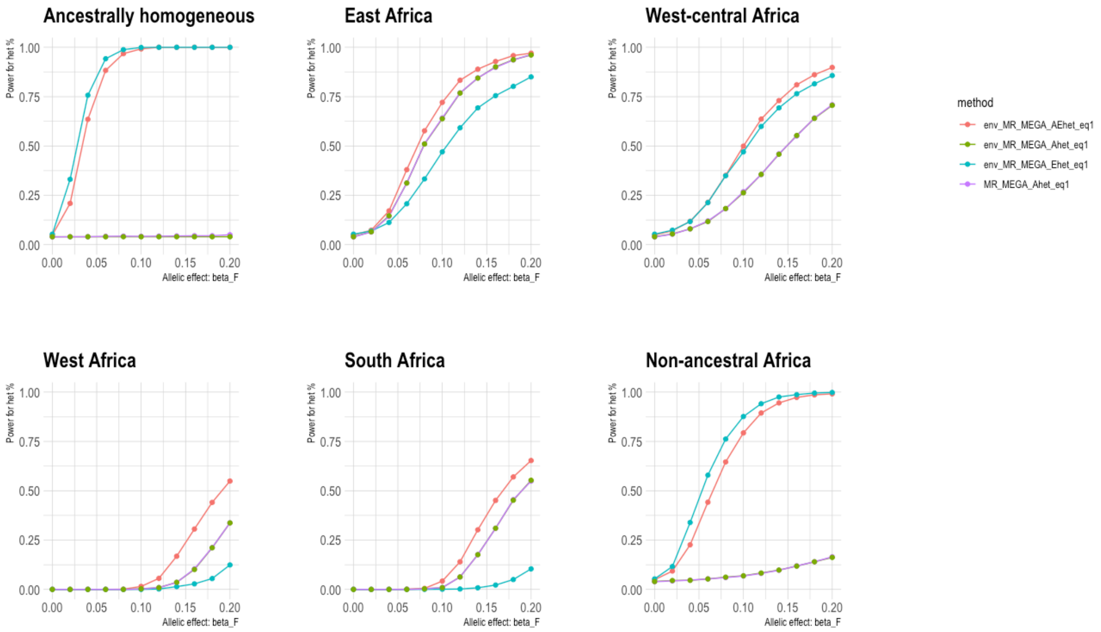

**Figure S4. Across six heterogeneity scenarios involving 16 sex-stratified cohorts, where female and male cohorts in the same population share the same smoking proportion (non-difference), env-MR-MEGA exhibits higher power in detecting association compared to MR-MEGA, especially in ancestrally homogeneous, west-central Africa and non-ancestral Africa scenarios.** "env_MR_MEGA_assoc" and "MR_MEGA_assoc" refer to the power to detect association obtained from env-MR-MEGA and MR-MEGA. Power was assessed at $P < 5 \times 10^{-8}$ and based on 1000 simulations with unequal sample sizes.

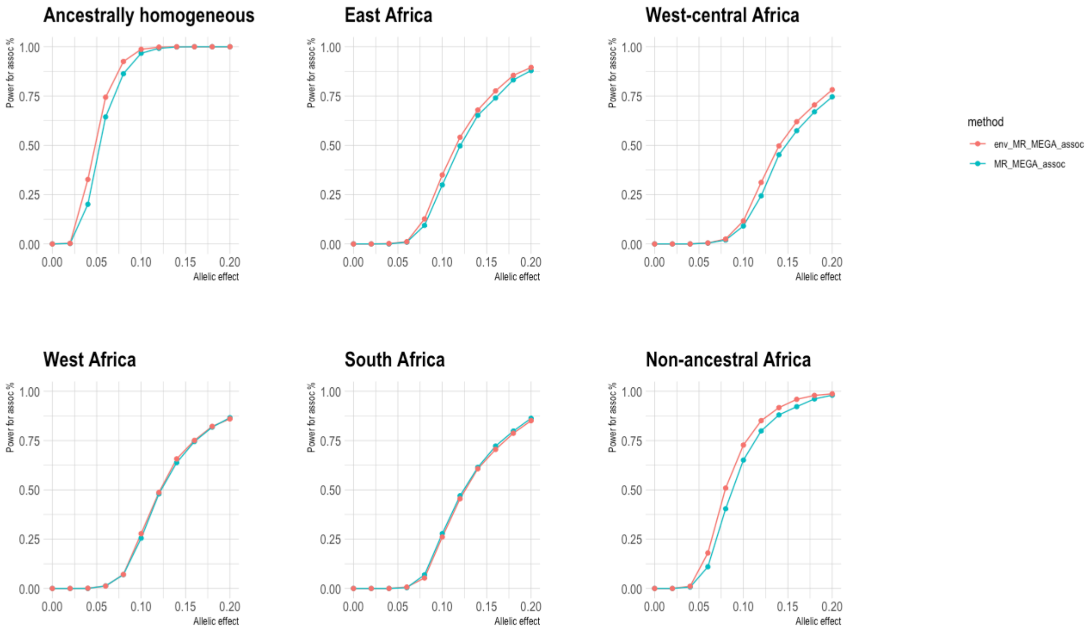

**Figure S5. Across six heterogeneity scenarios involving 16 sex-stratified cohorts, where minor reductions in smoking proportions between male and female cohorts occur (same direction), env-MR-MEGA exhibits higher power in detecting association compared to MR-MEGA, especially in ancestrally homogeneous, west-central Africa and non-ancestral Africa scenarios.** “env_MR_MEGA_assoc” and “MR_MEGA_assoc” refer to the power to detect association obtained from env-MR-MEGA and MR-MEGA. Power was assessed at $P < 5 \times 10^{-8}$ and based on 1000 simulations with unequal sample sizes.

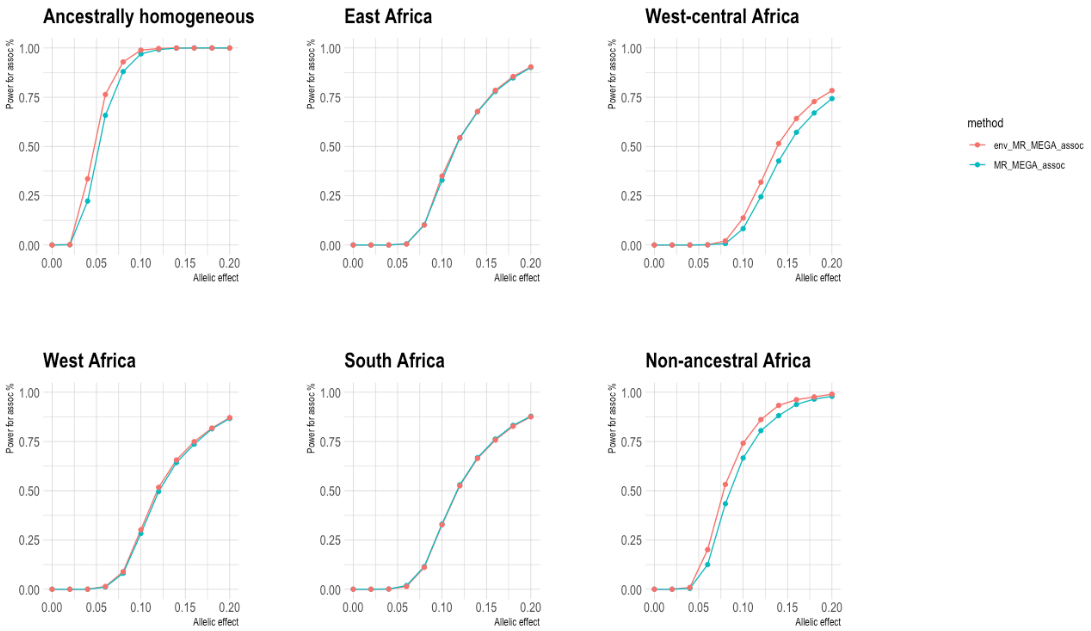

**Figure S6. When female and male cohorts in the same population share the same smoking proportion (no-difference) across six heterogeneity scenarios involving 16 sex-stratified cohorts, power for heterogeneity due to ancestry and environment from env-MR-MEGA exceeded power for heterogeneity due to ancestry from MR-MEGA, particularly in ancestrally homogeneity, west-central Africa and non-ancestral Africa scenarios.** "env_MR_MEGA_AEhet" (red line) corresponds to the power to detect heterogeneity due to ancestry and environment attained from env-MR-MEGA; "env_MR_MEGA_Ahet" (green line) corresponds to the power to detect heterogeneity due to ancestry alone attained from env-MR-MEGA; "env_MR_MEGA_Ehet" (blue line) corresponds to the power to detect heterogeneity due to environment alone attained from env-MR-MEGA; "MR_MEGA_Ehet" (purple line) corresponds to the power to detect heterogeneity due to ancestry attained from MR-MEGA. Power to detect heterogeneity due to ancestry and/or environment was assessed at $P < 0.05$.

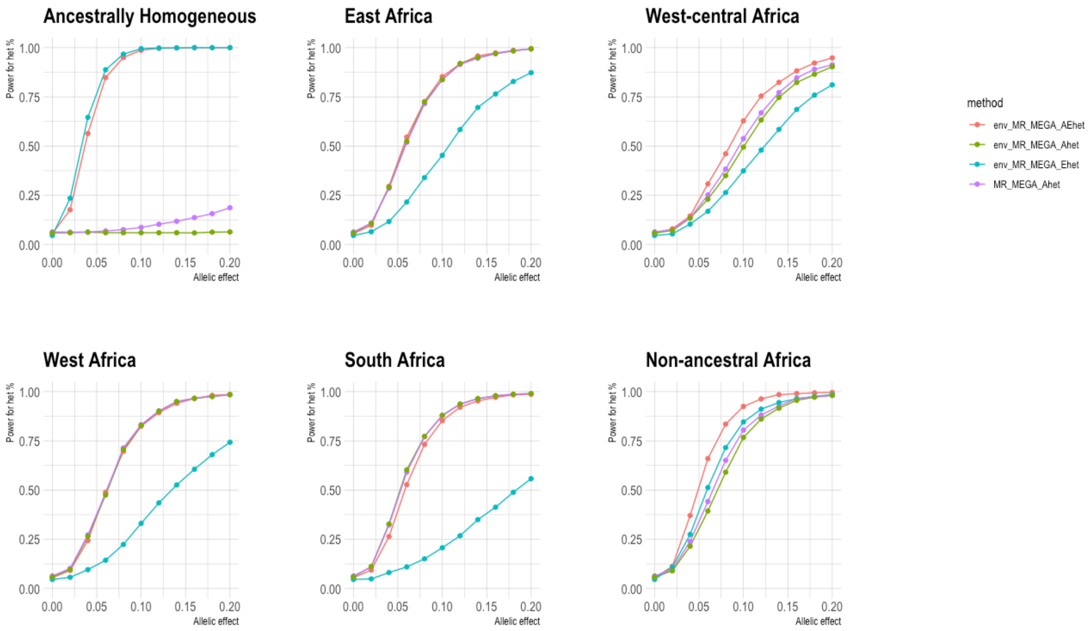

**Figure S7. When minor random reductions or increases in smoking proportions between male and female cohorts occur (mixed direction) across six heterogeneity scenarios involving 16 sex-stratified cohorts, power for heterogeneity in allelic and environmental effects from env-MR-MEGA exceeded power for heterogeneity in allelic effects from MR-MEGA, particularly in ancestrally homogeneity, west-central Africa and non-ancestral Africa scenarios.** “env_MR_MEGA_AEhet” (red line) corresponds to the power to detect heterogeneity due to ancestry and environment attained from env-MR-MEGA; “env_MR_MEGA_Ahet” (green line) corresponds to the power to detect heterogeneity due to ancestry alone attained from env-MR-MEGA; “env_MR_MEGA_Ehet” (blue line) corresponds to the power to detect heterogeneity due to environment alone attained from env-MR-MEGA; “MR_MEGA_Ehet” (purple line) corresponds to the power to detect heterogeneity due to ancestry attained from MR-MEGA. Power to detect heterogeneity due to ancestry and/or environment was assessed at $P < 0.05$.

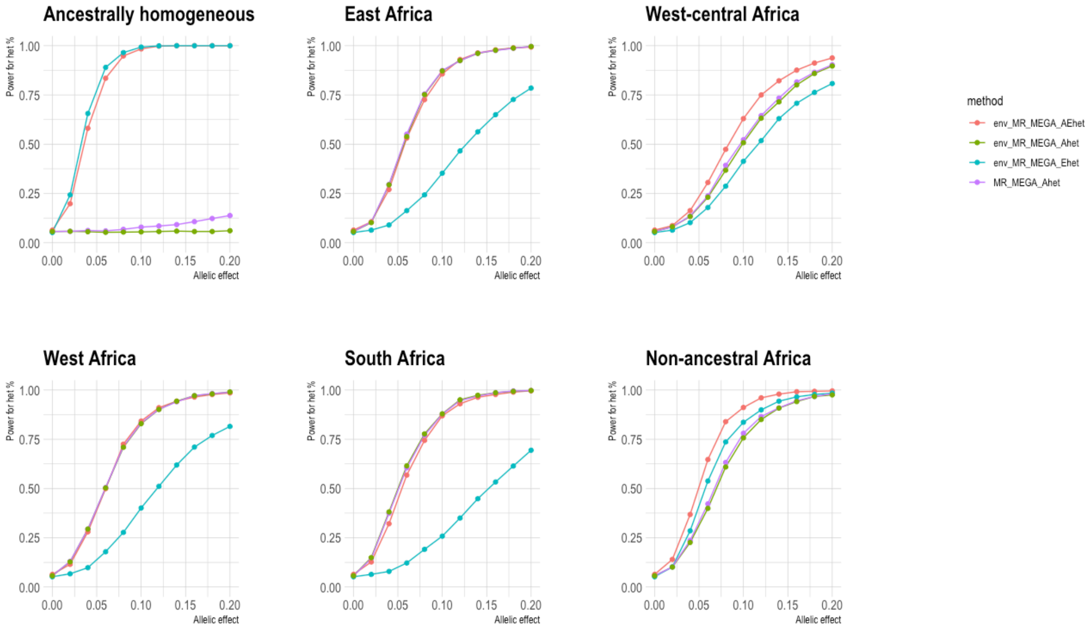

**Figure S8. When minor reductions in smoking proportions in female cohorts occur (same direction) across six heterogeneity scenarios involving 16 sex-stratified cohorts, power for heterogeneity due to ancestry and environment from env-MR-MEGA exceeded power for heterogeneity due to ancestry from MR-MEGA, particularly in ancestrally homogeneity, west-central Africa and non-ancestral Africa scenarios.** “env_MR_MEGA_AEhet” (red line) corresponds to the power to detect heterogeneity due to ancestry and environment attained from env-MR-MEGA; “env_MR_MEGA_Ahet” (green line) corresponds to the power to detect heterogeneity due to ancestry alone attained from env-MR-MEGA; “env_MR_MEGA_Ehet” (blue line) corresponds to the power to detect heterogeneity due to environment alone attained from env-MR-MEGA; “MR_MEGA_Ehet” (purple line) corresponds to the power to detect heterogeneity due to ancestry attained from MR-MEGA. Power to detect heterogeneity due to ancestry and environment was assessed at $P < 0.05$.

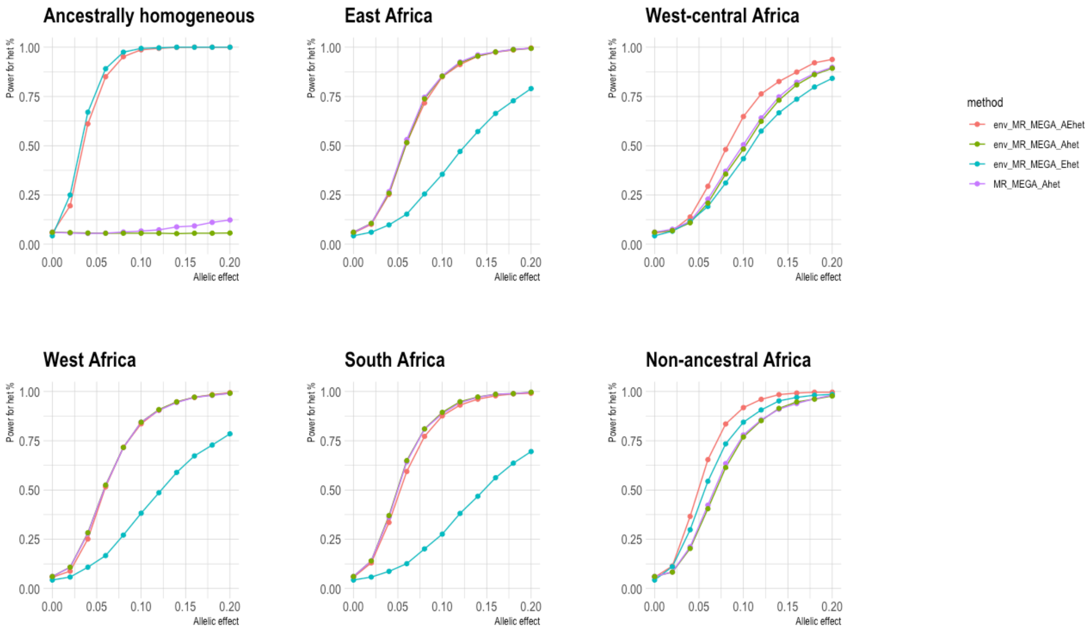

**Figure S9. When notable random reductions or increases in smoking proportions between male and female cohorts occur (mixed direction) across six heterogeneity scenarios involving 16 sex-stratified cohorts, power for heterogeneity due to ancestry and environment from env-MR-MEGA exceeded power for heterogeneity due to ancestry from MR-MEGA, particularly in ancestrally homogeneity, west-central Africa and non-ancestral Africa scenarios.** "env_MR_MEGA_AEhet" (red line) corresponds to the power to detect heterogeneity due to ancestry and environment attained from env-MR-MEGA; "env_MR_MEGA_Ahet" (green line) corresponds to the power to detect heterogeneity due to ancestry alone attained from env-MR-MEGA; "env_MR_MEGA_Ehet" (blue line) corresponds to the power to detect heterogeneity due to environment alone attained from env-MR-MEGA; "MR_MEGA_Ehet" (purple line) corresponds to the power to detect heterogeneity due to ancestry alone attained from MR-MEGA. Power to detect heterogeneity due to ancestry and/or environment was assessed at $P < 0.05$.

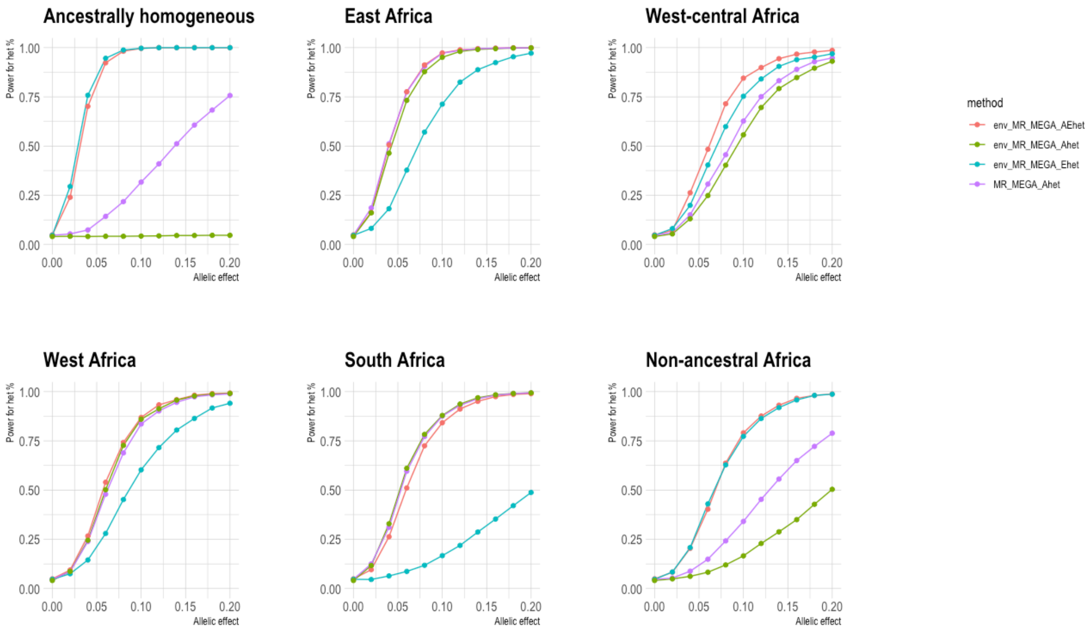

**Figure S10. When notable reductions in smoking proportions in female cohorts occur (same direction) across six heterogeneity scenarios involving 16 sex-stratified cohorts, power for heterogeneity due to ancestry and environment from env-MR-MEGA exceeded power for heterogeneity due to ancestry from MR-MEGA, particularly in ancestrally homogeneity, west-central Africa and non-ancestral Africa scenarios.**
"env_MR_MEGA_AEhet" (red line) corresponds to the power to detect heterogeneity due to ancestry and environment attained from env-MR-MEGA; "env_MR_MEGA_Ahet" (green line) corresponds to the power to detect heterogeneity due to ancestry alone attained from env-MR-MEGA; "env_MR_MEGA_Ehet" (blue line) corresponds to the power to detect heterogeneity due to environment alone attained from env-MR-MEGA; "MR_MEGA_Ehet" (purple line) corresponds to the power to detect heterogeneity due to ancestry attained from MR-MEGA. Power to detect heterogeneity due to ancestry and/or environment was assessed at $P < 0.05$.

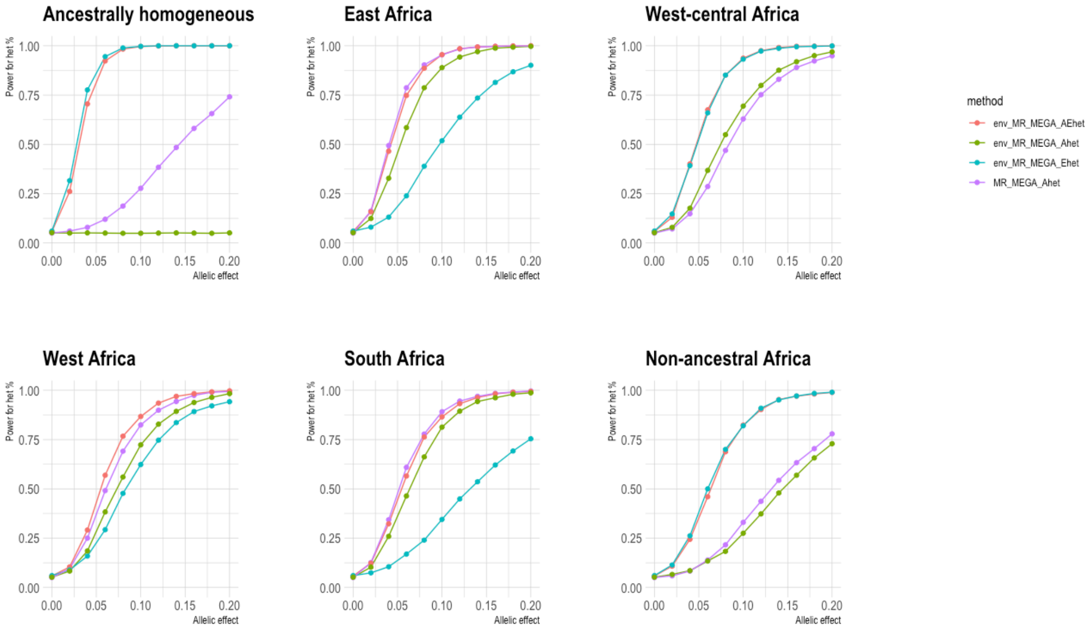
